# Sights and sounds of dolphins, *Tursiops truncatus,* preying on native fish of San Diego Bay and offshore in the Pacific Ocean

**DOI:** 10.1101/2022.03.02.482691

**Authors:** Sam H. Ridgway, Dianna S. Dibble, Mark Baird

## Abstract

For the first time, dolphins wearing video cameras were observed capturing and eating live native fish. While freely swimming in San Diego Bay, one dolphin caught 69 resident fish, 64 demersal, 5 near surface, while the other caught 40, 36 demersal and 4 near the surface. Two other dolphins were observed capturing 135 live native fish in a sea water pool. Two additional dolphins were observed feeding opportunistically during open water sessions in the Pacific Ocean. Notably, one of these dolphins was observed to consume 8 yellow-bellied sea snakes (*Hydrophis platurus*). Searching dolphins clicked at intervals of 20 to 50 ms. On approaching prey, click intervals shorten into a terminal buzz and then a squeal. Squeals were bursts of clicks that varied in duration, peak frequency, and amplitude. Squeals continued as the dolphin seized, manipulated and swallowed the prey. If fish escaped, the dolphin continued the chase and sonar clicks were heard less often than the continuous terminal buzz and squeal. During captures, the dolphins’ lips flared to reveal nearly all of the teeth. The throat expanded outward. Fish continued escape swimming even as they entered the dolphins’ mouth, yet the dolphin appeared to suck the fish right down.

## Introduction

Cameras have been attached to various marine mammals for observations of behavior under water (Davis *et al.* 2003). Monk seals have been observed capturing fish with these devices (Antonelis and Littnan 2008). Sounds and head jerks of toothed whales have been recorded as they captured live prey (Johnson *et al.* 2006; Jensen *et al.* 2009; Wisniewska *et al.* 2015, 2016). Sounds and video have also been recorded from dolphins finding and consuming dead fish (Wisniewska *et al.* 2014; Ridgway *et al.* 2014, 2015; Dibble *et al.* 2016; Ladegaard *et al.* 2017). However, sound and video together have never been used to observe behavior of dolphins and of the live fish they capture and consume. Our hypothesis was that we could make useful observations of dolphin’s fish captures employing inexpensive commercially available cameras that recorded video and audio. For the first time, two Navy dolphins wearing these cameras hunted and captured live fish while freely swimming in San Diego Bay. Two other Navy dolphins were observed capturing live fish in a sea water pool while two additional Navy dolphins were observed feeding opportunistically during open water sessions in the Pacific Ocean (Ridgway *et al.* 2018). Audio recordings from the camera allowed us to hear human audible sounds produced by the dolphins as they swam, searched, pursued, and captured prey. Video taken simultaneously by the same camera allowed us to begin relating dolphin sounds with behavior. Specifically we wanted to listen to dolphin sounds while they chase, capture and eat prey. We also wanted to see dolphin’s eye, throat and mouth movements during prey capture.

There are two main concepts on odontocete (toothed whales) feeding techniques. The first feeding concept involves strong intraoral suction which is used by most fish and marine mammals (Marshall and Golbogen 2015, Wainwright *et al.* 2015). For most odontocetes studied, observers have shown that suction for feeding is produced by hyolingual retraction and depression that withdraws the tongue and expands the throat to create a negative (sub-ambient) intraoral pressure. This negative intraoral pressure facilitates capturing, ingesting and manipulating prey in the mouth (Heyning and Mead 1989; Kastelein *et al.* 1997; Werth 2006*a*, *b*).

Ram feeding is a second concept (Marshall and Golbogen 2015, Wainwright *et al.* 2015). During ram feeding, prey is rapidly overtaken and clasped in jaws on the way to being swallowed. Bloodworth and Marshall (2005) summarize earlier assumptions about dolphin feeding when they state: “The stereotypical image of an odontocete is that of narrow, long snouted delphinids such as bottlenose dolphins (*Tursiops truncatus)*, which chase down prey with a clap-trap type of jaw containing numerous homodont teeth.” In studies comparing pygmy sperm whales (*Kogia spp.*) and *T. truncatus*, Bloodworth and Marshall (2005, 2007) observed that the former were mainly suction feeders while the later were mainly ram feeders. Feeding kinematic studies by Kane and Marshall (2009) demonstrated through direct measurement of pressure that several other species including *Delphinapterus leucas*, *Lagenorhynchus obliquidens* and *Globicephala melas,* used a combination of ram and suction feeding. Ito *et al.* (2002) compared *Tursiops aduncus* and *Neophocaena phocaenoides.* They suggested that these two species fed with a weak suction force compared to sperm whales (*Physeter macrocephalus*) and beaked whales (*Ziphius sp.*), which they regarded as strong suction feeders based on anatomy. No feeding on live fish was observed in these studies.

Marshall and Golbogen (2015) recommended video as an added tool to study marine mammal feeding. In our current study, we wanted to witness dolphin feeding strategies along with their concurrent sounds. With free-swimming dolphins searching for, pursuing, and capturing live prey that were actively trying to evade capture we though that we might be able to add to our knowledge about how dolphins find, capture and ingest prey.

## Materials and Methods

To observe dolphins catching live fish, we used cameras attached to the animals’ harness (Figure 1 A.) similar to our previous studies (Ridgway *et al.* 2015, 2018; Dibble *et al.* 2016). Our observations were of six *T. truncatus* of different ages (Table 1). four of the dolphins, B, K, S, Y, were collected from the wild from the Gulf of Mexico in the 1980s and obviously had experience in capturing live fish. Dolphin Z had been born at our facility in 2002 and Dolphin T had stranded on a Florida beach as a neonate with fresh umbilical in 2013 and rehabilitated at Sea World of Florida. Dolphins T and Z had never been observed to catch live fish prior to our observations.

**Figure 1.**
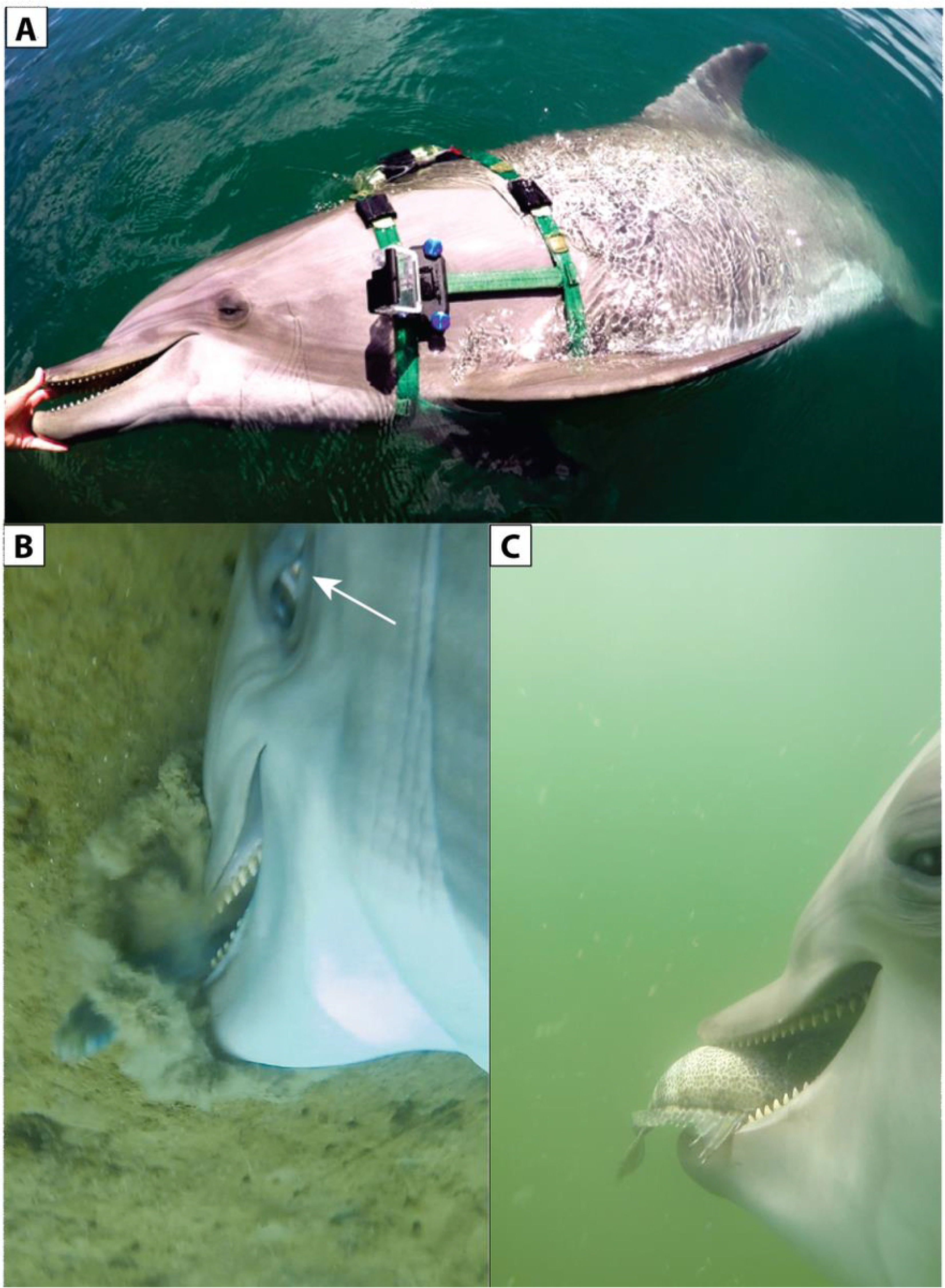
**A.** Dolphin S with camera attached to the left side of her harness. **B.** Dolphin S drills into the sea floor to seize a fish. Notice the white of the eye or sclera (arrow) shows the eye rotated toward the fish. **C.** Dolphin S brings out the fish with lips flaring in the posterior half of the gape area to show the upper tooth row and gular area expanding.

**Table1.** Navy dolphins observed

| **Dolphin** | **Sex** | **(cm)** | **(kg)** | **Age** | **DOB** | **Origin** | **Hrs** | **Sessions** | **Loc** |
| --- | --- | --- | --- | --- | --- | --- | --- | --- | --- |
| **S** | ♀ | **252** | **162** | **37** | **1980** | **MMA\*** | **14:00** | **15** | **SD Bay** |
| **K** | ♀ | **257** | **436** | **42** | **1977** | **MMA\*** | **4:00.00** | **5** | **SD Bay** |
| **B** | ♂ | **278** | **222** | **39** | **1978** | **MMA\*** | **1:30.00** | **2** | **Pool** |
| **T** | ♂ | **228** | **132** | **5** | **2012** | **Florida** | **1:30.00** | **2** | **Pool** |
| **Y** | ♀ | **246** | **172** | **34** | **1983** | **MMA\*** | **00:10** | **opportunistic** | **Ocean** |
| **Z** | ♀ | **259** | **176** | **15** | **2002** | **NMMP** | **00:04** | **opportunistic** | **Ocean** |

Table 1. Characteristics of the six bottlenose dolphins (*Tursiops truncatus*) used in the experiments. Dolphins S and K captured native fish while swimming freely in San Diego Bay. Dolphins B and T captured fish in a seawater pool. Dolphins Y and Z fished opportunistically while working in the Ocean. Dolphins S, K, B, T, and Y originated in the wild, Dolphin Z was born at the Navy facility. Dolphin T had stranded on a Florida beach as a very young animal, apparently still nursing. Dolphin T was rehabilitated by Sea World of Florida and determined by NOAA to be not releasable and was transferred to the Navy Marine Mammal Program in San Diego. We doubt that this animal had ever captured live fish. * MMA = Mississippi Management Area in the Gulf of Mexico off Gulfport, MS designated in the 1970s and 1980s for taking of dolphins permitted under the U. S. Marine Mammal Protection Act of 1972.

We recorded video and sound from four female and two male *T. truncatus* (Table 1) as they caught and ate live prey. All animals wore harnesses to which we attached cameras so that the dolphins’ fore bodies and the water ahead was clearly visible (Figure 1 A). The cameras were GoPro Hero3+, 4 or 4 session recording at 1080P at 60, 90 or 120 frames per second (FPS). The high FPS allowed us to observe changes in the dolphin’s behavior and body features frame by frame. The cameras recorded video and sound with a bandwidth of 16 kHz. This acoustic bandwidth is narrower than the usual dolphin clicks, which may reach 150 kHz or more (Au 1993). However, because the acoustic bandwidth of each click is so broad the camera system accurately records the timing of each click, pulse burst, squeal, or whistle. In previous studies we did simultaneous recordings with a broadband hydrophone system (125 kHz bandwidth) and the same cameras used here. In that study, the timing of the dolphin’s sounds were accurately represented by the camera even though recording a limited 16 kHz bandwidth (Ridgway *et al.* 2015, Dibble *et al.* 2016).

Dolphin S was recorded during 15 periods of about 50 minutes each. An observation period began when the dolphin fitted with harness and camera (Figure 1A.) was called from its netted enclosure to follow its trainer’s boat out to the bay. Out in the bay, the boat was stopped to allow the dolphin free rein to forage. After the 50-minute observation period, the dolphin followed the trainer’s boat back to its netted enclosure. These foraging periods occurred intermittently over 8 months resulting in 14 hours of video recordings.

Dolphin K was recorded during 5 periods of about 50 minutes each using the same procedure as for Dolphin S. These foraging periods occurred intermittently over 3 months resulting in 4 hours of video and sound recordings. In addition to the cameras, Dolphin K carried a sound trap that recorded sound at a 150 kHz bandwidth.

Two fish capture observation periods of 90 min each were recorded with two dolphins, B and T, in an above-ground 6 x 12 m sea water pool. Cameras were placed on both left and right sides of the harness so that the eye, the total gape area, the rostrum and the external gular area were clearly visible as the animals swam. Live Pacific mackerel (*Scomber diego)*, Pacific Sardines *(Sardinops sagax caerulea)* and Northern Anchovies (*Engraulis mordax)* were acquired from a local San Diego Bay live bait supplier about 500 m from the pool where the dolphins swam. The live fish were transported in an aeriated sea water container and were introduced to the pool so the dolphin’s feeding and simultaneous sounds could be observed.

Incidental prey captures were observed for two dolphins (Y and Z) freely swimming in the open ocean (Ridgway *et al.* 2018). These two dolphins wore cameras located dorsally behind the blowhole. The blowhole and rostrum was visible, but not the mouth and teeth. We could listen to their sounds and observe their head and rostrum movements as they took prey during 4 minutes of prey encounters by Z and 10 minutes of small fish encounters by Y.

To identify and classify sounds, recordings were displayed in Adobe Audition CS6 with Blackmann-Harris windows of 512 points. The recordings were then inspected aurally and visually. Video, still pictures and sequences were clipped and correlated in real time with the sounds dolphins produced as they searched for and captured prey allowing the anatomical functions present before, during and after prey capture to be observed.

### Ethics Statement

The study followed protocols approved by the Animal Safety Committee of the U.S. Navy Marine Mammal Program and followed all animal care guidelines of the Institutional Animal Care and Use Committee at the Naval Information Warfare Center Pacific and the Navy Bureau of Medicine and Surgery, and followed all applicable U.S. Department of Defense guidelines for the care and use of animals.

## Results

### Observations of Dolphins S and K foraging in San Diego Bay

Dolphins foraged in San Diego Bay over an area of about 2 square Km. Video recorded 69 fish captures by S (64 demersal fish and 5 near the surface) and 40 fish captures (36 demersal and 4 near the surface) by K. Prey were identified by appearance in the video and comparison with published sources (https://www.wildlife.ca.gov/Fishing/Ocean/Fish-ID/Sportfish). Identified prey included spotted sand bass (*Paralabrax maculatofasciatus*), barred sand bass (*Paralabrax nebulifer*), top smelt (*Atherinops affinis*), yellowfin croaker (*Umbrina roncador*), California halibut (*Paralichthys californicus*) and pipefish (*Syngnathus leptorhynchus*). Four predation events included either corvina (*Cilus gilberti*) or white sea bass (*Atractoscion nobilis*), but fish identification could not be confirmed. In all cases, dolphins S and K pursued one fish at a time. There were 4 instances when rays were seen resting on the sea floor, but S did not attempt to capture these.

Once prey was captured (typically by the tail), dolphin S usually opened her mouth quite wide to reorient the fish (head first) for ingestion. It seemed to us that the fish would be able to escape during this time. However, as S opens her mouth it is very noticeable that the gular region is expanded, tooth rows are exposed and the upper and lower lips flare out even when the dolphins is backing out of a bottom capture (Figures 1B,1C and 2A). From these views, it appears that the tongue is withdrawn, the gular muscles expand the throat, and in addition the lips flare as the fish is sucked in. As S closes her mouth again with the fish secured, the video reveals water and debris coming out of her mouth (video Figure S1). During the capture sequence, it does not appear that the teeth ever fully penetrate the fish or that the dolphin’s jaws are fully closed.

**Figure 2.**
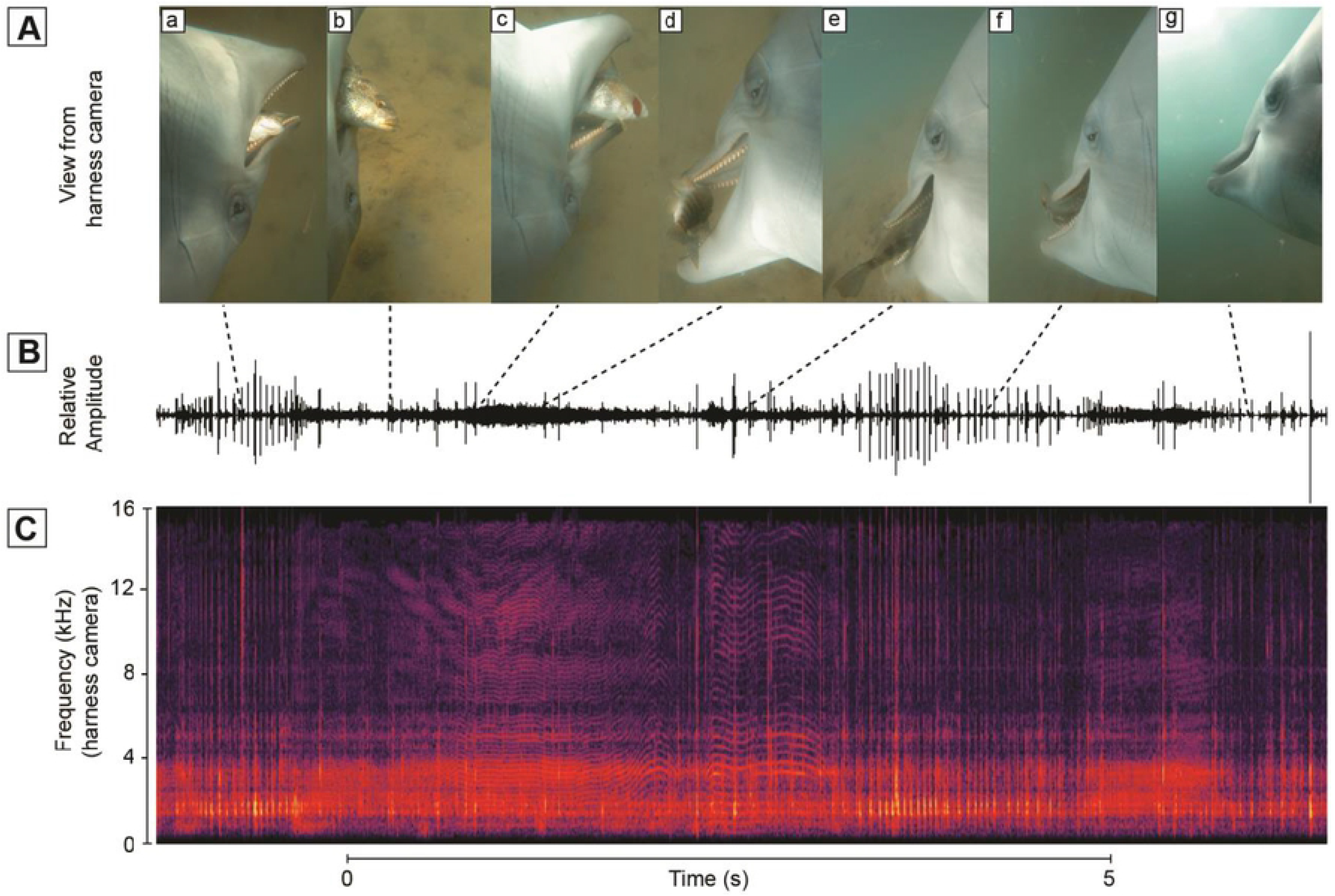
**A**. View of dolphin fore body while capturing fish. a. the dolphin contacts the fish, b. the fish flees as the eye focuses on the fish, c. the fish escapes, d. is found again. e. and is swallowed **B.** Relative amplitude of sound recorded as dolphin S located, chased and captured wild fish. **C**. Spectrogram of audible sound showing variations in pulse rate and peak frequency characteristic of a squeal.

**Figure S1**. Video of Dolphin S capturing fish in San Diego Bay

As dolphins hunted, they clicked almost constantly at intervals of 20 to 50 ms. On approaching prey, click intervals shorten into a terminal buzz and then a squeal (Ridgway *et al.* 2015). On contact with fish, buzzing and squealing was almost constant until after the fish was swallowed (Figures 2 and 3). It became apparent during video analyses when dolphins had identified the next prey target. The background noise would quickly intensify masking many dolphins sounds as the animal picked up speed in pursuit of a fish. Often times, after a few fish chases, the dolphin’s heartbeat could also be heard on the video.

**Figure 3.**
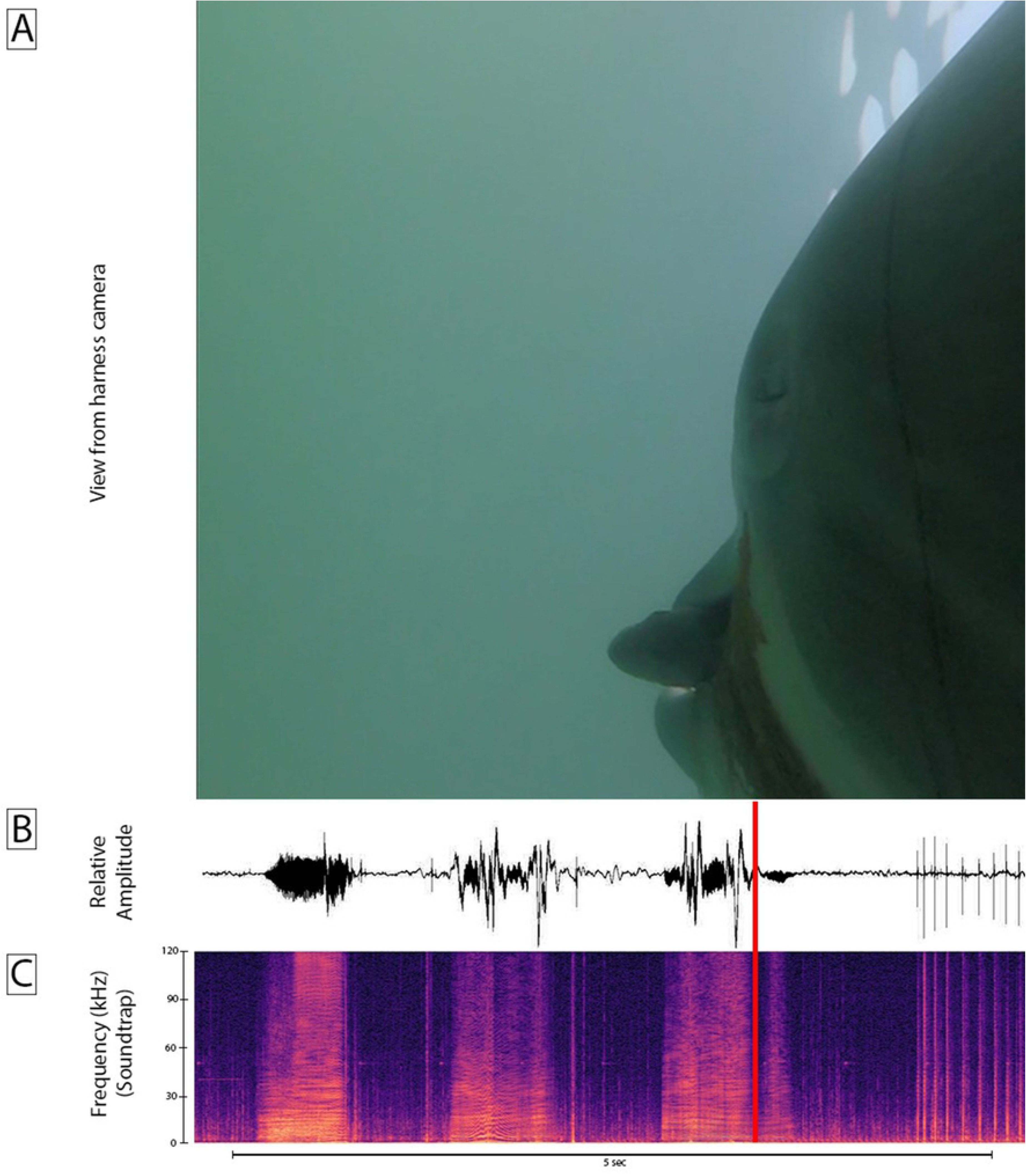
Dolphin K captures live fish in San Diego Bay. **A.** View of dolphin K while capturing fish. **B.** Relative amplitude of sound recorded from dolphin K during the capture observed in **A**. **C**. Audible frequencies of sound recorded from dolphin K during the capture observed in **A**. The red bar shows the point of capture. Echolocation clicks appear at the end of this recordings as the dolphin searches for another fish

As the dolphins approached prey, it was evident that the visible eye was oriented toward the fish. The white posterior sclera was a visible sign that the eye ball rotated forward toward prey. At times we see a ring of skin deformation surrounding the eye ball that is probably indicative of eye muscle contractions. Further, when the dolphin pursued fish near the surface the animal oriented its body ventrally toward the surface and the visible eye rotated toward the surface as well (Figure 4A). Three times as S pursued prey, the fishes circled back towards the dolphin. If the fishes escaped predation and swam past the dolphin, we observed the anterior white sclera as the dolphin rotated the eyeball backward to track the fish as it fled (Figure 4B).

**Figure 4.**
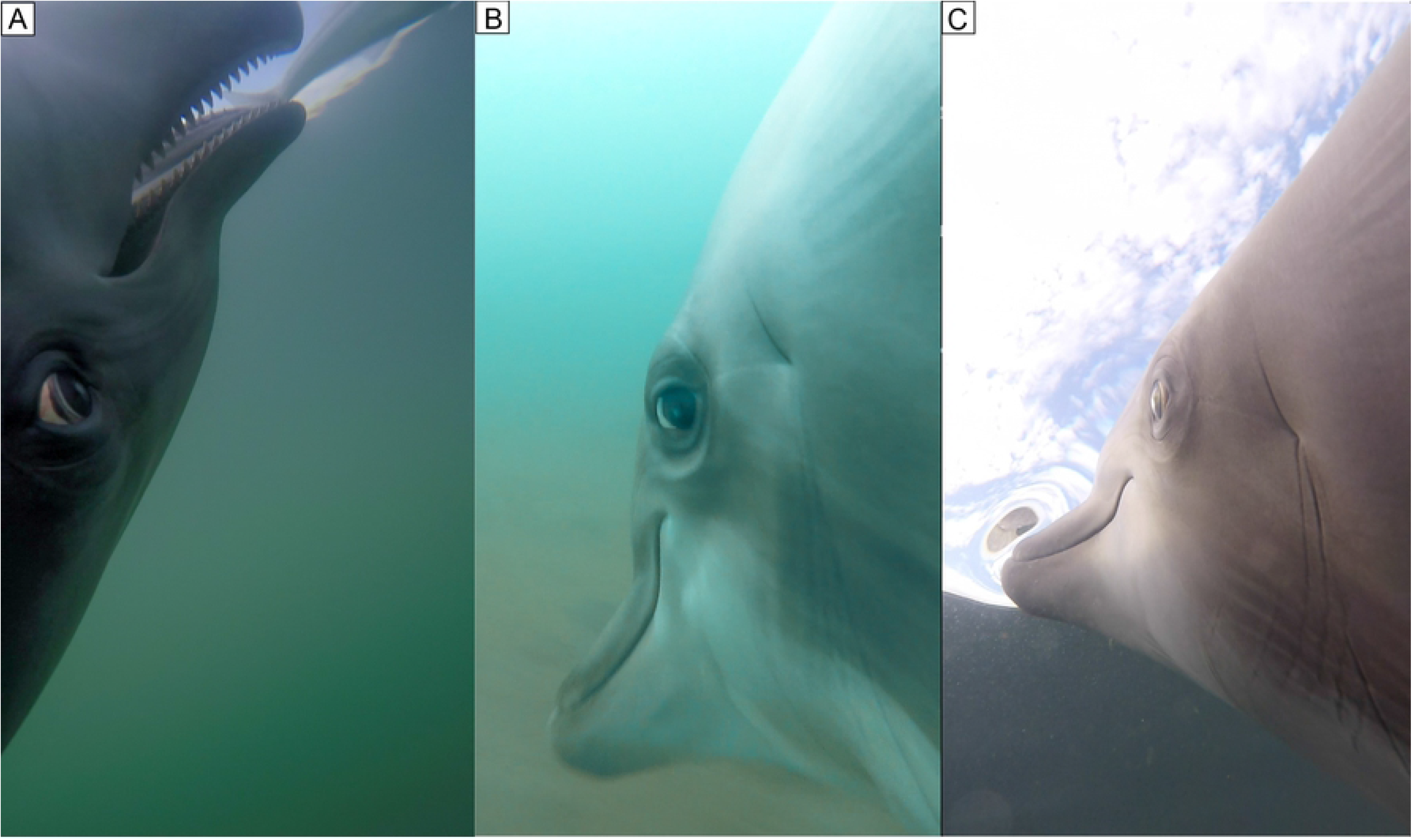
Dolphin S visually tracks fish by rotating eye. **A**. During a surface chase, the dolphin visually tracks a fish by turning upside down and by rotating the eye forward displaying the posterior sclera. **B**. The dolphin rotates the eye backward displaying the anterior sclera as a fish escapes to the rear of the dolphin. Note the depression under the eyeball attending the backward rotation. **C**. Dolphin S eye without a depression showing.

Several times, dolphin S was observed chasing a smelt, *A. affinis,* near the surface (Figure 4A). With the dolphin in pursuit, the smelt would often jump out of the water traveling a meter or more. In these events, the dolphin swam ventral up while tracking the prey and met the fish as it landed back in the water. In contrast, the majority of *P. maculatofasciatus* and the few *P. nebulifer* captured were located on the sea floor, most often in patches of vegetation. Bass (*P. maculatofasciatus)* were often observed resting on their pectoral fins on the sea floor. As the dolphin approaches, the bass scurries to hide in a patch of bottom vegetation. During this time, S is observed buzzing and squealing to find the hidden fish, then she dives into the vegetation to bring out a mouth full of bottom sediment, fish, and plant material (video Figure S1). The fish is swallowed and the plant material and sediment are ejected back into the water. Dolphin S is able to secure and manipulate the fish in her mouth while riding her mouth of the vegetation. This process resulted in the fish escaping her jaws two times. During one of these examples, the fish was recaptured; the other time, the fish evaded capture.

### Observations of Dolphins B and T foraging in a clear seawater pool

We observed three hours of live fish captures by dolphin’s B and T in a 6 × 12 m sea water pool. The two dolphins were placed in the pool at the same time. Live Pacific mackerel (*Scomber diego)*, Pacific Sardines *(Sardinops sagax caerulea)* and Northern Anchovies (*Engraulis mordax)* were acquired from a local San Diego Bay live bait supplier about 500 m from the pool. The live fish where rapidly transported and placed in pool where the dolphins swam. Dolphin B, a 39-year-old (Table 1) taken from the Gulf of Mexico in the 1980s, began catching fish almost immediately.

Cameras were placed on both left and right sides of the harness so that the eyes, the full gape area, the rostrum and the external gular area were clearly visible as the animals swam. Our plan was to observe both right and left eyes during these sessions. However, the dolphins seized the fish with a swipe of the head to one side or the other. This side swipe motion allowed the eye to be viewed only from the camera on one side. The older dolphin B (Table 1) immediately began to prey on the fish, sucking them in with a side swipe of his head (Figure 4A). At first T just swam in the midst of the schooling fish without consuming them. Perhaps after noticing the predatory behavior of B, dolphin T began to take fish. His captures were attended by much squealing (Figure 5 B and C). During the 3 hours of recording, dolphin B took 114 fish while T only consumed 21.

**Figure 5.**
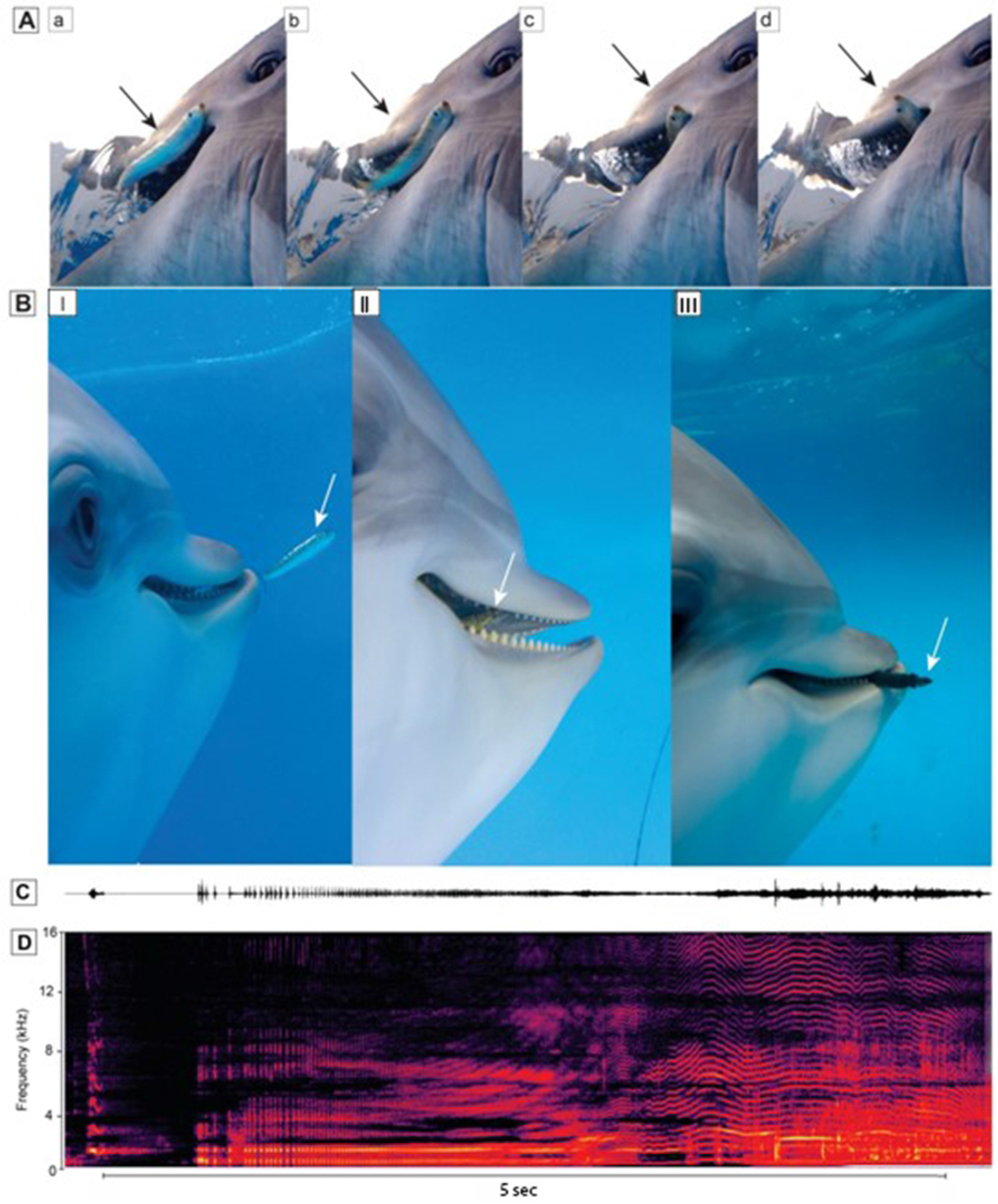
Dolphins capture live fish in a seawater pool. **A**. Dolphin B sequence (a, b, c, d) shows B sucking in a live fish (arrow) in his mouth. **B**. Dolphin T (a) locates a fish, right eye rotated forward. (b) Upon capture, the lower posterior lip is pulled down showing the gums and teeth and fish (arrow) inside the mouth. (c) The dolphin reorients the fish while still pulling the lip down and expanding the gular area apparently eliciting intraoral pressure reduction, yet the fish almost escapes. **C.** Relative amplitude of sound recorded from dolphin T during the capture observed in **B**. **D**. Audible frequencies of sound recorded from dolphin T during the capture observed in **B**.

Consistent with previous observations of Dolphin S, the visible eye rotated toward the fish. It appeared that Dolphin B was actively sucking the fish into his mouth as fish swimming movements were observed as the fish entered the mouth near the lip commissure (Figure 4A).

### Observations of incidental captures by Dolphins Z and Y in the open ocean

Several incidental captures were noted from dolphin’s Z and Y during an unrelated objective as they searched the open waters of the Pacific Ocean for mine simulators (Ridgway *et al.* 2018). In these cases, we noticed the rostrum of the dolphin move quickly to the side as the prey disappeared near the lip commissure. These captures were attended by an audible squeal by the dolphin (Ridgway *et al.* 2015). In most cases the prey appeared to be small fish that we could not identify. It is notable that on one day, dolphin Z preyed on 8 yellow bellied sea snakes (*Hydrophis platurus*). The dolphin clicked as it approached the snake and then sucked it in with a bit more heard jerking as the flopping snake tail disappeared and the dolphin made a long squeal (Figure 6, and Video Figure S2). A larger snake successfully fled (Video Figure S3).

**Figure 6.**
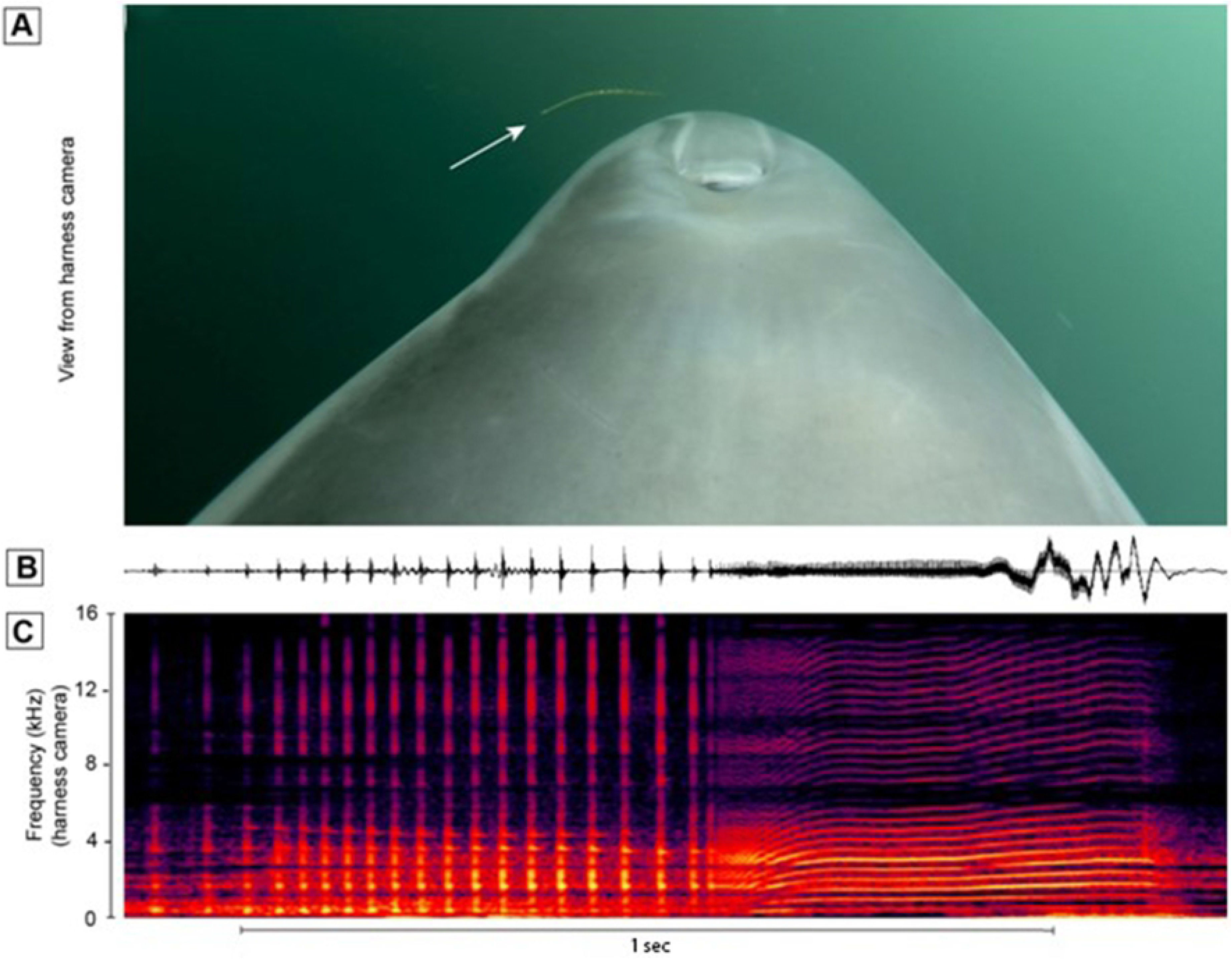
**A**. View of dolphin fore body while nearing sea snake **B.** Amplitude of sound recorded as dolphin Z located and captured a sea snake. Head jerks (abrupt amplitude changes at the end of **B**) are indicative of swallowing prey (c.f. Ridgway et al., 2015) **C**. Spectrogram of audible sound.

**Figure S2**. Video Dolphin Z catching sea snakes in the Pacific Ocean with obvious head jerks and a victory squeal.

**Figure S3.** Video Slowed down video of an escaping sea snake with notable dolphin clicks

## Discussion

Even in a few hours of hunting in San Diego Bay, dolphins S and K found and consumed fish from the bottom and just under the surface. This is consistent with the wide variety of prey know from this species in the wild. Concerning the prey of bottlenose dolphins, Wells and Scott (1999) state, “This extensive variety of prey inhabits an equally diverse selection of habitats, and includes benthic-reef and sandy-bottom prey and their associated predators, pelagic schooling fish and cephalopods, and deeper-water fish.” The yellow bellied sea snake (*Hydrophis platurus)* (Graham *et al.* 1987) (Figure 6, Video Figures S 2 and S3) is an outlier. We were at first skeptical that our dolphin had preyed on sea snakes and searched for possible fish of similar appearance. We were able to rule out other possibilities due to appearance of the head, tail, body marking and swimming motions of the snakes (Video Figures S 2 and S3). Furthermore, near the time of our observed predation, some yellow-bellied sea snakes stranded on a beach within about 2 km of the site of our observations. Dolphin Z that consumed sea snakes was aged 15 and had been born at our facility. Perhaps the dolphin’s lack of experience in feeding with dolphin groups in the wild led to the consumption of this outlier prey.

In our observations of Dolphins S, K, B and T, where an eye was visible in the video, it appeared that the dolphin always had the eye rotated toward the fish. When a fish jumped into the air the dolphin swam, ventrum up, and the eye tracked the fish through the air. When the fish reentered the water, the dolphin was there to catch it. This upside down feeding near the surface has been observed previously in wild *T. truncatus* by Leatherwood (1975). When a fish swam past the dolphin the eye rotated to track the fish as it fled behind the dolphin. These dolphins appeared to use both and sight and sound to find prey.

Wood and Evans (1980) suggested that bottlenose dolphins can track the precise movements of prey at close range by listening for the prey’s swimming sounds. But dolphins may also use passive listening to detect prey over longer distances. Most prey eaten by bottlenose dolphins are soniferous (Barros and O’Dell 1990, Gannon *et al.* 2005). These fish produce sounds that are primarily below 5 kHz, but many have calls that contain some energy above 10 kHz, well within the hearing range of bottlenose dolphins and within the sensitivity range of our camera microphones. We did not hear fish sounds in our recordings nor can we comment on detection of swimming motion by our dolphins. The observations of Wood and Evans (1980) were from the very quiet environment of a redwood pool. Our observations with cameras worn by the swimming dolphins have a higher level of background noise. Certainly, dolphins may listen for fish sounds. Our observations all contained sonar pulses and squeals by the dolphins.

We were surprised by the frequency with which Dolphins S and K caught fish on the bottom of San Diego Bay. In fact, the majority of captures were from the bottom. The dolphins readily dove into bottom vegetation and even into the bottom sediment to capture fish. On sandy bottoms, *T. truncatus* has been observed to dive into the bottom sand up to their pectoral flippers to acquire fish (Rossbach and Herzing 1997). In San Diego Bay we did not observe Dolphins going that far into the bottom to acquire fish. The majority of captures were of demersal fish which appears to be a common type of prey for *T. truncatus* in other areas (Barros and O’Dell 1990, Wells and Scott 1999, Blanco *et al.* 2001).

We were surprised by the ability of all of our dolphins to open their upper and lower lips near the commissure to allow prey to be sucked in to the mouth. It has been observed that this area has a high density of nerve endings and the greatest skin sensitivity (Ridgway and Carder 1993). With years of experience in feeding dolphins, we had not noticed this this lip motion until we held the rostrum to limit mouth opening to 2 cm or so and saw that small fish could be sucked in at the commissure. The perspective from the camera just behind the dolphin’s mouth helped to view this lip action. The placement of the camera on the side of the dolphin enabled us to view the lifting of the lip to reveal the complete row of the dolphin’s teeth (Figures 1 B and C; 5B). Harper *et al.* (2008) note that the lateral rostral muscles insert into the lip, which is a dense and immobile connective tissue structure in delphinids; the lateral rostral muscles also insert into the lateral dense connective tissue surrounding the melon. Our observations show that the posterior lips served by the lateral rostral muscle move or flair to take in fish. The lower lip also was pulled down to reveal the gum and lower tooth row. The muscles that support this action do not extend far up the lower jaw (Figure 8). To our knowledge this muscular action has not been previously observed.

During the chase, the dolphins that had cameras placed laterally on their side, (B, K, S, T) could be seen quickly opening their mouths, exposing the teeth and gums by flaring the lips and noticeably expanding their gular regions. Perhaps these were fish capture attempts or a preparatory phase to suction.

Some of our observations of fish captures near the surface might be described as ram feeding. However, the great majority feeding events appeared to rely mainly on suction. Rather than seizing fish in a “claptrap” of the toothy beak, dolphins appeared to mostly suck in fish from the side with lips opened, gular area expanded, and tongue withdrawn to increase intraoral space creating negative pressure. Bottlenose dolphins have a robust hyoid apparatus with strongly developed gular musculature to enable strong suction.

### Structures that expand the throat, depress the tongue and open the lips at the commissure

After our observations of dolphins feeding, we reviewed literature on hyoid bones and gular musculature of *T. truncatus* (Lawrence and Schevill,1965; Heyning and Mead, 1989; Reidenberg and Laitman, 1994; Werth, 2007). We wanted to consider the lip opening at the commissure and gular expansion and apparent suction that we had observed (Figures 1, B and C., Figure 2 and 4). We had available a reconstructed skeleton that enabled us to label the hyoids (Figure 7). Also, we had available some MRI and CT scans from previous studies of living dolphins (Ridgway et al., 2006) to show the muscles, hyoids and adjacent structures in situ (Figure 7). Also, from a previous study (Ridgway, 1999) we had sections from a *T. truncatus* that died of natural causes. From these we were able to view muscles that serve to move upper and lower lips near the commissure (Figure 8).

**Figure 7.**
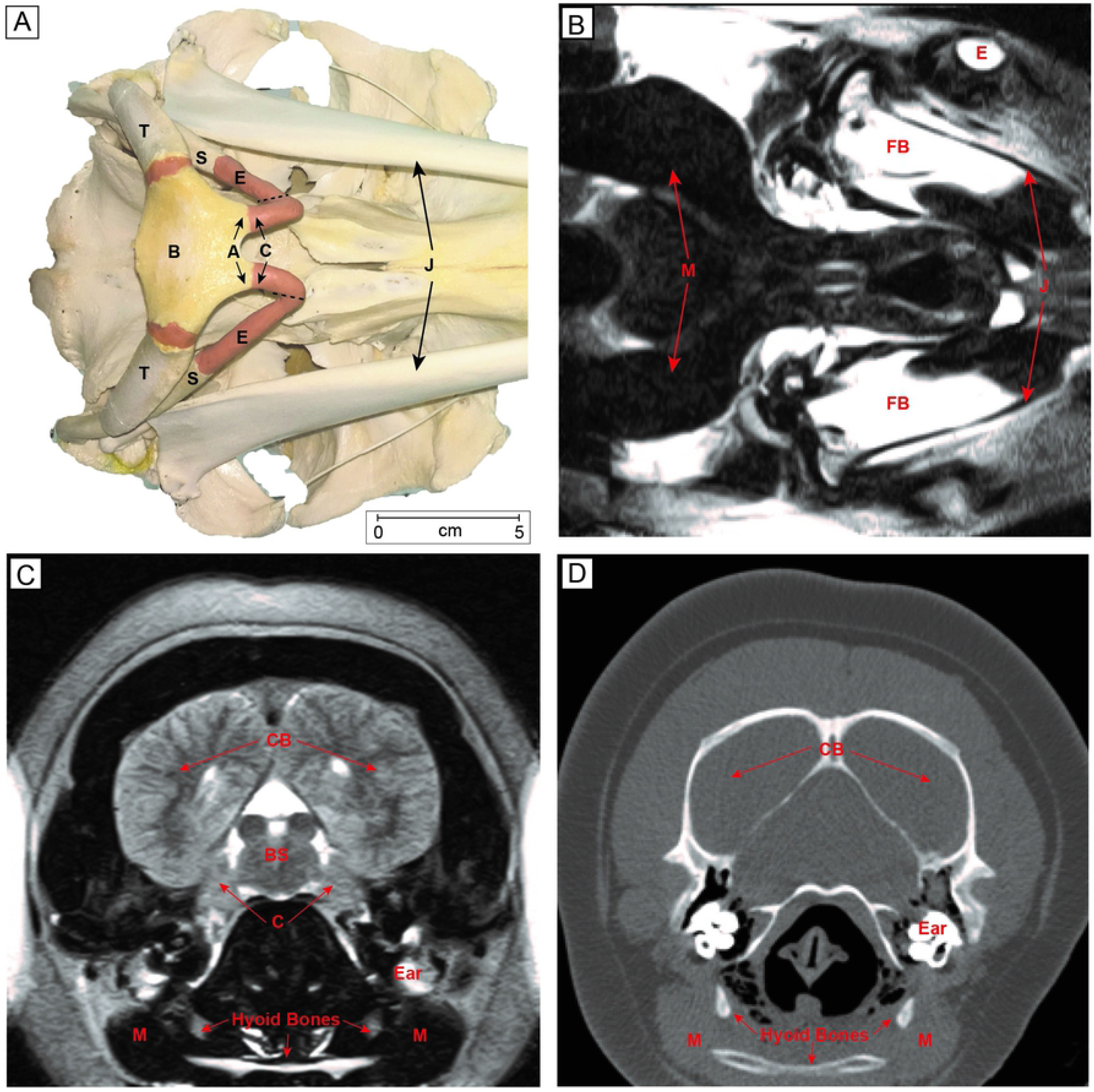
Bones and muscles related to the hyolingual apparatus bottlenose dolphins, *Tursiops truncatus.* **A.** Ventral view of a dolphin skull. A, articular process; B, basihyal; C, ceratohyal; E, epihyal; S, stylohyal; T, tympanohyal; J, mandible. Hyoid bone terminology from Reidenberg and Laitman (1994); **B.** Horizontal MRI image viewed at the level of the stylohyal showing the heavy (dark) musculature. FB, fat body of lower jaw, E eye, J jaw. **C.** Frontal MRI image displaying anterior portions of brain and hyoid apparatus. C, cerebellum; CB, cerebrum; BS, brainstem, Ear, M, muscle masses of hyoid area. **D.** Frontal CT scan shows hyoid bones and immense musculature adjacent M. (MRI and CT scans are reconfigured and relabeled from Ridgway *et al.* 2006).

**Figure 8.**
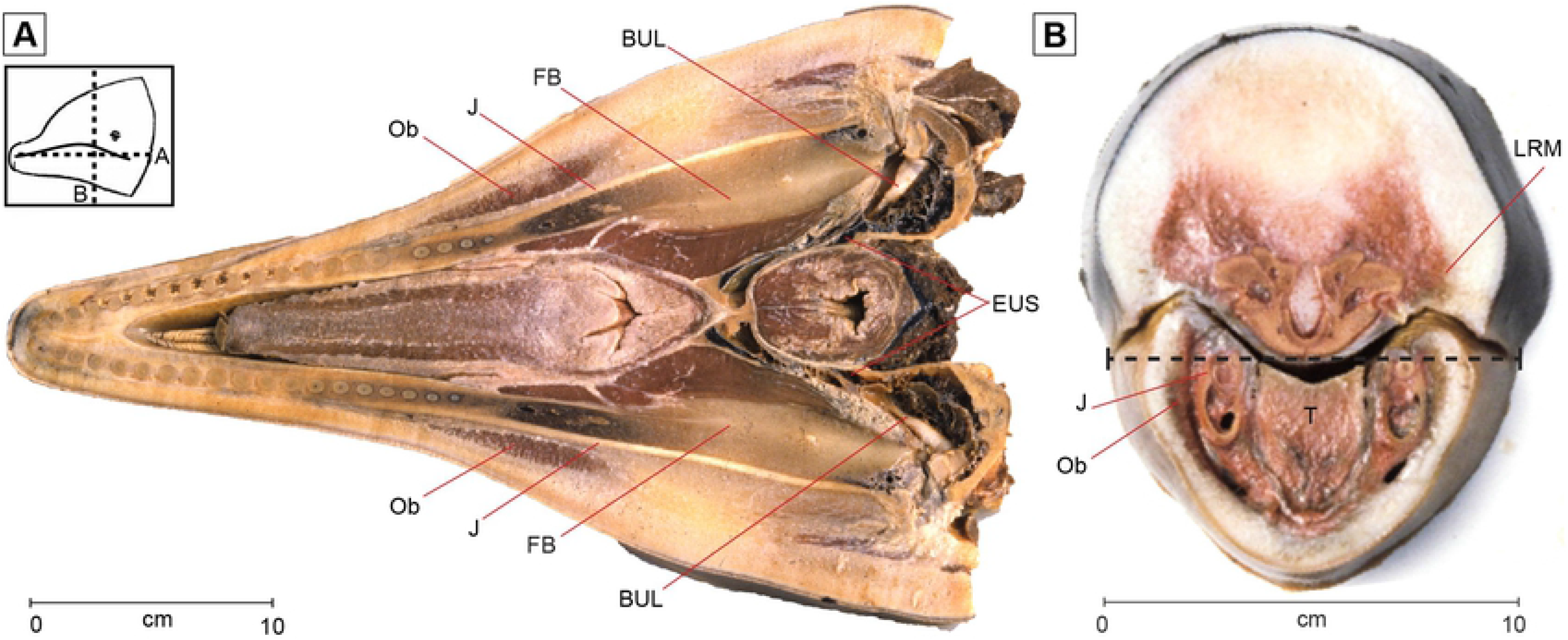
Two sections through the dolphin mouth showing relationships of some muscles of the upper and lower lip. The box shows the orientations of **A**. and **B**. **A**. horizontal section through lower jaw. Ob orbicularis oris, muscles of lower lip, J Jaw, FB, fat body of jaw, BUL auditory bulla or tympanic bone, EUS Eustachean tube. **B** Frontal section through the mouth just anterior to the lip commissure. Dashed line shows the approximate orientation of **A**. Ob orbicularis oris muscles of lower lip, LRM lateral rostral muscle. J, jaw bone, T, tongue.

## Acknowledgments

We thank Pixie Rixon Whaling and Elaine Allen who trained and worked with dolphin S employed in our study. Jaime Kennemer was the trainer for dolphins Z and Y. Her efforts in supporting our project to observe trained dolphins in the open ocean is much appreciated. The study would not have been possible without their expert training. We also thank the supervisors and trainers who assisted with dolphins B and T. We thank Alison E. Cramer and Cheryl Short for painstakingly putting together the skeleton of a female *T. truncatus* that died at our facility. This specimen was used for showing the dolphin hyoid.

## Literature Cited

Antonelis, G. and C. Littnan. Monk seal research and recovery effort: how Crittercam revolutionized our understanding of monk seal ecology. In G. Marshall, ed. Animal borne imaging symposium. National Geographic, Washington D.C. 2008:15–20

Au, W. W. L. The sonar of dolphins. 1993. Springer-Verlag, New York, NY.

Barros, N. B. and D. K. O’Dell. Food habits of bottlenose dolphins in the southeastern United States. In S. Leatherwood and R. R. Reeves, eds. Academic Press, San Diego, CA. 1990:309–28

Blanco, C., Salomón, O., and J. A. Raga. 2001. Diet of the bottlenose dolphin (*Tursiops truncatus*) in the western Mediterranean Sea. Journal of the Marine Biological Association of the United Kingdom 2001 81:1053–1058.

Bloodworth, B. and C. D. Marshall. 2005. Feeding kinematics of *Kogia* and *Tursiops* (Odontoceti: Cetacea): characterization of suction and ram feeding. Journal of Experimental Biology 208:3721–3730.

Bloodworth, B. E. and C. D. Marshall. A functional comparison of the hyolingual complex in pygmy and dwarf sperm whales (*Kogia breviceps* and *K. sima*), and bottlenose dolphins (*Tursiops truncatus*). Journal of Anatomy 2007 211:78–91.

Davis, R. W., Fuiman, L. A., Williams, T. M., Horning, M. and W. Hagey. Classification of Weddell seal dives based on 3 dimensional movements and video-recorded observations. Marine Ecology Progress Series 2003 264:109–122.

Dibble, D. S., Van Alstyne, K. R. and S. Ridgway. Dolphins signal success by producing a victory squeal. International Journal of Comparative Psychology 2016 29:1.

Gannon, D. P., Barros, N. B., Nowacek, D. P., Read, A. J., Waples, D. M. and R. S. Wells. Prey detection by bottlenose dolphins, *Tursiops truncatus*: an experimental test of the passive listening hypothesis. Animal Behaviour 2005 69:709–720.

Graham, J. B., Lowell., W. R., Rubinoff, I. and J. Motta. Surface and subsurface swimming of the sea snake *Pelamis platurus*. Journal of Experimental Biology 1987 127:27–44.

Harper, C. J., Mclellan, W. A., Rommel, S. A., Gay, D. M., Dillaman, R. M. and D. A. Pabst. Morphology of the melon and its tendinous connections to the facial muscles in bottlenose dolphins (*Tursiops truncatus*). Journal of Morphology 2008 269:820–839.

Heyning, J. E. and J. G. Mead. Contributions in science. Natural History Museum of Los Angeles County 1989 405:1–64.

Ito, H., Ueda, K., Aida, K. and T. Sakai. Suction feeding mechanisms of the dolphins. Fisheries science 2002 68:268–271.

Jensen, F. H., Bejder, L., Wahlberg, M. and P. T. Madsen. Biosonar adjustments to target range of echolocating bottlenose dolphins (*Tursiops sp.*) in the wild. Journal of Experimental Biology. 2009 212:1078–1086.

Johnson, M., Madsen, P. T., Zimmer, W. M. X., de Soto, N. A. and P. L. Tyack. Foraging Blainville’s beaked whales (*Mesoplodon densirostris*) produce distinct click types matched to different phases of echolocation. Journal of Experimental Biology 2006 209:5038–5050.

Kane, E. A. and C. D. Marshall. Comparative feeding kinematics and performance of odontocetes: belugas, Pacific white-sided dolphins and long-finned pilot whales. Journal of Experimental Biology 2009 212:3939–3950.

Kastelein, R.A., Staal, C., Terlouw, A. and M. Muller. Pressure changes in the mouth of a feeding harbour porpoise (*Phocoena phocoena*). In A. J. Read, P. R. Wiepkma, and P. E. Nachtigall, eds. The biology of the harbor porpoise. DeSpil Publishers, Woerden, Netherlands. 1997:279–291.

Beedholm, K., da Silva, V. M. F. and P. T. Madsen. 2017. Ladegaard, M., Jensen, F. H., Amazon river dolphins (*Inia geoffrensis*) modify biosonar output level and directivity during prey interception in the wild. Journal of Experimental Biology: 2017 220:2654–65

Lawrence, B. and W.E. Schevill. Gular musculature in delphinids. Bulletin of the Museum of Comparative Zoology 1965 133:1–65.

Leatherwood, S. Some observations of feeding behavior of bottle-nosed dolphins (*Tursiops truncatus*) in the northern Gulf of Mexico and (*Tursiops* cf. *T. gilli*) off southern California, Baja California, and Nayarit, Mexico. Marine Fisheries Review 1975 37:10–16.

Marshall, C. D. and J. A. Goldbogen. Feeding mechanisms. In M. A. Castellini and J. A. Mellish, eds. Marine mammal physiology: requisites for ocean living. CRC Press, Boca Raton, FL. 215:95–118

Reidenberg, J. S. and J. T. Laitman. Anatomy of the hyoid apparatus in odontoceli (toothed whales): specializations of their skeleton and musculature compared with those of terrestrial mammals. The Anatomical Record 1994 240:598–624.

Ridgway, S.H. and D.A. Carder. Features of dolphin skin with potential hydrodynamic importance. IEEE Engineering in Medicine and Biology 1993 12:83–88.

Ridgway, S.H. 1999. An illustration of Norris’ acoustic window. Marine Mammal Science 15:926–930.

Ridgway, S. H., Houser, D., Finneran, J. J., Carder, D. A., Keogh, M., Van Bonn, W., Smith, C. R., Scadeng, M., Mattrey, R. and C. Hoh. Functional imaging of dolphin brain metabolism and blood flow. Journal of Experimental Biology 2006 209:2902–2910.

Ridgway, S. H., Moore, P. W., Carder, D. A. and T. A. Romano. Forward shift of feeding buzz components of dolphins and belugas during associative learning reveals a likely connection to reward expectation, pleasure and brain dopamine activation. Journal of Experimental Biology 2014 217:2910–2919.

Ridgway, S., Dibble, D. S., Van Alstyne, K. and D. Price. On doing two things at once: dolphin brain and nose coordinate sonar clicks, buzzes and emotional squeals with social sounds during fish capture. Journal of Experimental Biology 2015 218:3987–3995.

Ridgway, S. H., Dibble, D. S. and J. A. Kennemer. Timing and context of dolphin clicks during and after mine simulator detection and marking in the open ocean. Biology Open 2018 7:bio031625.

Rossbach, K. A. and D. L. Herzing. Underwater observations of benthic feeding bottlenose dolphins (*Tursiops truncatus*) near Grand Bahama Island, Bahamas. Marine Mammal Science 1997 13:498–504.

Wainwright, P. C., McGee, M. D., Longo, S. J. and L. Patricia Hernandez. Origins, innovations, and diversification of suction feeding in vertebrates. Integrative and Comparative Biology 2015 55:134–145.

Wells, R. S. and M. D. Scott. Bottlenose dolphin *Tursiops truncatus* (Montagu,1821). In S. H. Ridgway and R. J. Harrison, eds. Handbook of marine mammals: the second book of dolphins and porpoises, vol 6. Academic Press, San Diego, CA. 1999:137–182

Werth, A. J. Mandibular and dental variation and the evolution of suction feeding in Odontoceti. Journal of Mammalogy 2006a 87:579–588.

Werth, A. J. Odontocete suction feeding: experimental analysis of water flow and head shape. Journal of Morphology 2006b 267:1415–1428.

Werth, A. J. Adaptations of the cetacean hyolingual apparatus for aquatic feeding and thermoregulation. Anatomical Record 2007 290:546–568.

Wisniewska, D. M., Johnson, M., Nachtigall, P. E. and P. T. Madsen. Buzzing during biosonar-based interception of prey in the delphinids *Tursiops truncatus* and *Pseudorca crassidens*. Journal of Experimental Biology 2014 217:4279–4282.

Wisniewska, D. M., Ratcliffe, J. M., Beedholm, K., Christensen, C. B., Johnson, M., Koblitz, J. C., Wahlberg, M. and P. T. Madsen. Range-dependent flexibility in the acoustic field of view of echolocating porpoises (*Phocoena phocoena*). 2015 E-life 4: e05651.

Wisniewska, D. M., Johnson, M., Teilmann, J., Rojano-Doñate, L., Shearer, J., Sveegaard, S., Miller, L. A., Siebert, U. and P. T. Madsen. Ultra-high foraging rates of harbor porpoises make them vulnerable to anthropogenic disturbance. Current Biology 2016 26:1441–1446.

Wood, F. G. and W. E. Evans. Adaptiveness and ecology of echolocation in toothed whales. In R. G. Busnel and J. F. Fish, eds. Animal sonar systems. Plenum, New York, NY. 1980:381–426.

